# Rapid wing size evolution in African fig flies (*Zaprionus indianus*) following temperate colonization

**DOI:** 10.1101/2024.11.15.623845

**Authors:** Weston J. Gray, Logan M. Rakes, Christine Cole, Ansleigh Gunter, Guanting He, Samantha Morgan, Camille R. Walsh-Antzak, Jillian A. Yates, Priscilla A. Erickson

## Abstract

Invasive species often encounter novel selective pressures in their invaded range, and understanding their potential for rapid evolution is critical for developing effective management strategies. *Zaprionus indianus* is an invasive drosophilid native to Africa that reached Florida in 2005 and likely re-establishes temperate North American populations each year. We addressed two evolutionary questions in this system: first, do populations evolve phenotypic changes in the generations immediately following colonization of temperate environments? Second, does *Z. indianus* evolve directional phenotypic changes along a latitudinal cline? We established isofemale lines from wild collections across space and time and measured twelve ecologically relevant phenotypes, using a reference population as a control. *Z. indianus* evolved smaller wings following colonization, suggesting early colonizers have larger wings, but smaller wings are favorable after colonization. No other phenotypes changed significantly following colonization or across latitudes, but we did see significant post-colonization changes in principal components of all phenotypes. We documented substantial laboratory evolution and effects of the laboratory environment across multiple phenotypes, emphasizing the importance of controlling for both possibilities when conducting common garden studies. Our results demonstrate the potential for rapid adaptation in *Z. indianus*, which could contribute to its success and expansion throughout invaded ecosystems.

## Introduction

Invasive and introduced species often adapt rapidly to their new environments, facilitating their spread and potentially exacerbating their ability to harm the native ecosystem, agriculture, or human health (Prentis et al. 2008; Whitney and Gabler 2008; Borden and Flory 2021). Many documented cases of evolution in invasive organisms come from invasions that are decades or even centuries old. For example, California poppies, which were introduced to Chile in the nineteenth century, are larger in the invasive range (Leger and Rice 2003). Cane toads, introduced to Australia in 1935, have evolved skeletal changes that facilitate more efficient dispersal, accelerating the rate of range expansion (Phillips et al. 2006; Hudson et al. 2016). Marine copepods have repeatedly evolved physiological tolerance to fresh water in multiple invasions over the past 200 years (Lee 1999). Less is known about evolution in the immediate aftermath of an invasion, though some studies have documented evolution of invasive species on shorter time scales. For example, less than 20 years after colonization, invasive populations of speckled wood butterflies showed differences in body size and wing morphology in a common garden experiment (Hill et al. 1999). However, relatively few studies have tested for rapid evolution in the generations immediately following colonization of a new environment. Understanding the potential for adaptation early in invasions would enhance our understanding of rapid evolution in natural populations and might inform management strategies to mitigate the impacts of invasive species.

The rapid establishment of geographic clines in the invasive range can reflect adaptation to local environmental conditions that vary across the cline. For example, *Medicago polymorpha* has evolved a latitudinal flowering time cline in North America that mirrors the cline found in its native range in Europe (Helliwell et al. 2018). The fruit fly *Drosophila subobscura* evolved clines in wing morphology and body size within 20 years of colonizing North America, and both clines match those found in the native range (Huey et al. 2000; Gilchrist et al. 2001). Thus, environmental gradients across latitudes can create selective pressures that lead to rapid local adaptation and differentiation within the invasive range. Latitudinal clines for a variety of fitness-related and morphological traits are described in both North America and Australia in natural populations of the model organism *D. melanogaster* (Flatt 2020). *D. melanogaster* was introduced to both continents within the last several hundred years (David and Capy 1988), so these studies provide a template for the types of traits that might evolve rapidly in other introduced insects.

The African fig fly, *Zaprionus indianus* (Gupta), is a unique drosophilid model for studying invasion biology because it likely re-invades temperate habitats across a latitudinal gradient in North America each year. *Z. indianus* was originally introduced to the western hemisphere in Brazil in 1999 (Vilela 1999) and then spread northwards into central and North America, reaching Florida by 2005 (Linde et al. 2006). It was first detected in Virginia in 2012 and has reached as far north as Minnesota, Quebec, and Ontario (Renkema et al. 2013; Holle et al. 2018). Although *Z. indianus* primarily feeds on a wide variety of decaying fruits (Lachaise et al. 1988), it is a pest of figs (EFSA Panel on Plant Health (PLH) et al. 2022) and potentially a pest of soft-skinned fruits like raspberries (Pfeiffer et al. 2019; Allori Stazzonelli et al. 2023). *Z. indianus* appears to lack cold tolerance (Pfeiffer et al. 2019): it is first detected later in the summer than related drosophilids, and it disappears earlier in the fall (Rakes et al. 2023). Sub-freezing temperatures likely cause local extirpation of the population each year, followed by re-colonization the following year. In some locations, *Z. indianus* is found in one year but not the next, suggesting these populations are not permanently established and recolonization is idiosyncratic (Holle et al. 2018; Gleason et al. 2019; Rakes et al. 2023). *Z. indianus* reaches sexual maturity approximately 16 days after egg laying (Nava et al. 2007), suggesting that several (4-5) generations likely occur between its establishment in temperate locations in July and its extirpation in November (Rakes et al. 2023). Furthermore, despite an initial bottleneck, *Z. indianus* populations in eastern North America maintain surprisingly high levels of genetic diversity (Comeault et al. 2020, 2021) and a wide thermal niche breadth (Comeault et al. 2020). This high level of genetic diversity combined with a short generation time raises the possibility that *Z. indianus* might evolve rapidly following recolonization of temperate environments. Adaptation could potentially fuel further expansion of the species or establishment of permanent populations; therefore, understanding the mechanisms and extent of adaptation is important to informing its management as a potential pest.

Other drosophilids undergo rapid evolution in response to seasonal changes over short time scales, informing the potential for rapid evolution in *Z. indianus*. In temperate environments, fly populations undergo several generations of reproduction as the seasons change from spring to summer to fall, and the changing environment imposes selection that influences the genetic makeup of the population (Bergland et al. 2014). This selection alters fitness-related phenotypes, reflecting the classical life history tradeoff between survival and reproduction. Populations in the summer adapt to high resource availability and favorable conditions for population growth, whereas overwintering populations have traits that favor survival over reproduction. For example, in *D. melanogaster*, stress tolerance, reproductive traits, and immune function all vary seasonally (Miyo et al. 2000; Schmidt and Conde 2006; Dev et al. 2013; Behrman et al. 2015, 2018). Wing morphology also changes throughout the growing season (Tantawy 1964; Önder and Aksoy 2022). Based on these findings, we hypothesized that morphological, stress tolerance, and life history traits might also evolve over the course of a single growing season as *Z. indianus* responds to temperate environments following invasion.

Common garden experiments are a standard approach to test for genetically determined phenotypic variation. One challenge for common gardens, especially those conducted on samples separated in time, is to mitigate any potential effects of adaptation of the study organism to the laboratory environment (Hoffmann and Ross 2018). For example, *D. melanogaster* rapidly evolves reduced stress resistance when reared in laboratory culture (Hoffmann et al. 2001). A second potential complication is unintended variation in assay conditions (Moloney et al. 2009; Suckow and Tirado-Muñiz 2023). For example, seasonal changes in cold hardening and cold tolerance in field-caught populations *of D. melanogaster* also occurred in a lab-reared control population, suggesting the variation observed was due to unintentional differences in rearing or assay conditions (Stone et al. 2020). Two possible experimental strategies are generally proposed: controlling and minimizing the number of generations in the lab (resulting in samples being assayed at different times with potentially different assay conditions) (Behrman et al. 2015; Rudman et al. 2022), or assaying all individuals at the same time under common assay conditions (but after a different number of generations in the lab) (Ueno et al. 2023). Both approaches have the potential to capture variation that does not reflect true biological variation of the original populations. Therefore, using carefully controlled studies that account for possible adaptation to lab culture and/or potential variation in assay conditions is essential to accurately document temporal phenotypic changes in natural populations.

In this study, we used a common-garden approach to compare morphological and life history traits between wild-derived *Z. indianus* collected across space (a latitudinal gradient in eastern North America) and time (a single temperate growing season). We collected flies early and late in the growing season from two orchards in Virginia in 2022 and from four North American locations spread across ∼15° latitude. Using carefully controlled laboratory experiments and a reference population, we specifically sought to test three hypotheses:

1. *Z. indianus* undergoes rapid phenotypic evolution following colonization of temperate environments
2. *Z. indianus* exhibits local adaptation of phenotypes along a cline stretching from Florida to Connecticut
3. *Z. indianus* evolves rapidly under laboratory culture conditions

Our results suggest that *Z. indianus* evolves rapidly in the field but also adapts rapidly in laboratory culture, suggesting a high evolutionary potential in this invasive species with pest potential. Our findings also highlight the importance of rigorous controls in temporal experiments due to unintended variation in laboratory conditions that can influence phenotypes.

## Materials and Methods

### Experimental design and timeline

We tested for post-colonization (early vs late season) and spatial (latitudinal) variation in *Z. indianus* phenotypes by collecting wild flies from orchards, rearing them in the lab as isofemale lines for 3-4 generations, and comparing phenotypes across seasons and locations (Figure 1). Because the early and late flies were captured and assayed approximately three months apart from one another, we also wanted to control for potential variation in the lab environment or assay conditions that could impact our phenotypic measurements. We included a highly inbred line as a control that was phenotyped alongside every batch (Olazcuaga et al. 2022). This line should be genetically uniform and phenotypically consistent across generations; any changes in the phenotype of this line over time are likely phenotypically plastic responses to variation in the laboratory environment or assay variability.

**Figure 1:**
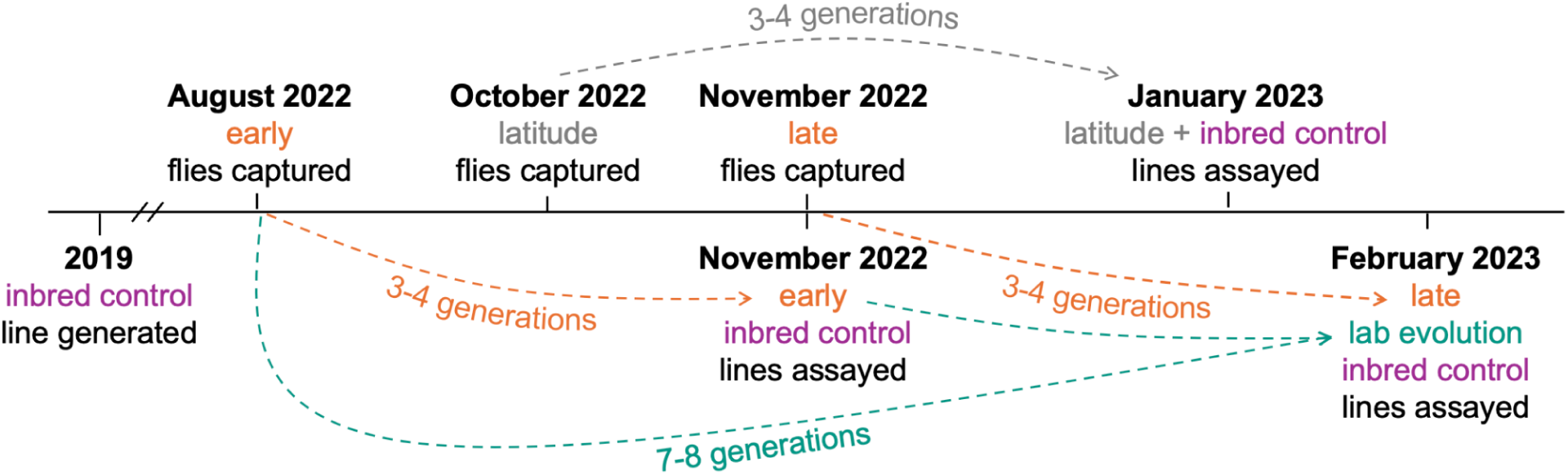
Overview of experimental design and timeline. The diagram indicates the relative timing of fly capture and experimental assays for the post-colonization (orange), latitude (gray), and lab evolution (green) experiments. The post-colonization evolution experiment compares the phenotypes of isofemale lines captured early and late in the season. The lab evolution experiment compares phenotypes of lines captured early in the season after 3-4 generations of lab rearing and 7-8 generations of lab rearing. The latitude experiment compares lines collected from locations spanning 15° latitude in North America. The inbred control line (purple) serves as a reference to test for lab environment or assay variation between experiments conducted at different times.

Isofemale lines are not genetically uniform and have the potential to evolve as they adapt to laboratory conditions (David et al. 2005). To measure potential laboratory evolution in *Z. indianus*, we randomly chose twelve lines (six from each Virginia orchard) collected in the early season and re-phenotyped them alongside the lines collected late in the season (Figure 1, green). Thus, the re-phenotyped lines were kept in the lab for an additional ∼4 generations for a total of ∼7-8 generations. Comparing the second round of phenotyping to the first round of phenotyping allowed us to assess potential laboratory evolution.

### Control line

The control line was derived from flies originally collected in Charlottesville, VA in August 2019. The line was inbred with full sibling matings for 10 generations and then reared in the lab with one generation every 3-4 weeks until the beginning of our experiments in August 2022. Each major set of phenotyping (early, late, and latitude) was separated into two batches which were completed within 14 days of each other; controls were included in each batch.

### Fly collections and isofemale line generation

All flies were collected by netting or aspirating off of fallen fruit from orchards in the fall of 2022 (Table 1; see Rakes et al. 2023 for additional collection details). For the post-colonization evolution experiment, flies were collected in August and November from two orchards in Virginia, approximately 100 km from each other. For the latitudinal assay, flies were collected from four locations spanning the eastern United States from Florida to Connecticut. Upon returning to the lab, we held wild-caught flies in bottles containing 50 mL cornmeal-molasses medium, a slice of banana, and live yeast for 24 hours to encourage mating. We placed individual females in vials containing 10 mL cornmeal-molasses media sprinkled with live yeast. After 7 days, we removed the females and the offspring were reared to adulthood. We recorded the proportion of females that successfully produced offspring for each collection as the isofemale success rate. All flies were reared in an incubator at 27°C, 50% relative humidity, 14L:10D light cycle.

**Table 1:** Fly collection details. Isofemale line success rate is the proportion of wild-caught females that produced at least one offspring in the lab. The number of isofemale lines phenotyped is the number of lines measured in the pupation time assay; some lines were not measured for all other phenotypes due to variation in the number of adult flies obtained. See Rakes *et al*. (2023) for additional information about collections.

| Location | Latitude, longitude | Collection Date | Isofemale line success rate (total) | Number of isofemale lines phenotyped |
| --- | --- | --- | --- | --- |
| Carter Mountain Orchard, Charlottesville, VA | 37.991, -78.472 | 8/19/2022 (early) | 0.88 (117) | 30 |
|  |  | 11/10/2022 (late) | 0.29 (100) | 19 |
| Hanover Peach Orchard, Mechanicsville, VA | 37.572, -77.266 | 8/17/2022 (early) | 0.84 (128) | 27 |
|  |  | 11/9/2022 (late) | 0.63 (100) | 29 |
| Fruit and Spice Park, Homestead, FL | 25.525, -80.493 | 10/2/2022 | 0.81 (200) | 18 |
| Hillcrest Orchard, Ellijay, GA | 34.620, -84.373 | 9/30/2022 | 0.76 (80) | 15 |
| Linvilla Orchard, Media, PA | 39.885, -75.410 | 10/14/2022 | 0.66 (80) | 17 |
| Lyman Orchard, Middlefield, CT | 41.494, -72.730 | 10/13/2022 | 0.55 (80) | 16 |

### Rearing controlled-density vials

After 1-2 generations in the lab, we expanded each isofemale line in bottles containing 50 mL media. Prior to collecting eggs for the phenotyping generation, adults were held in fresh bottles containing fly media, a banana slice, and live yeast for ∼3 days to encourage mating. We then constructed laying chambers using inverted plastic fly bottles with air holes and a petri dish containing 3% agar and 30% grape juice concentrate. We added ∼200 µL of live yeast paste to the agar and allowed flies to oviposit for 24 hours. After removing adults, the petri dishes were held for 24 additional hours to allow larvae to hatch. We used a piece of flattened wire to gently extract ∼30 first-instar larvae and placed them in a fresh food vial with 10 mL media. Five vials were collected for each line. We reared at least 12 vials of the control line for each batch, and control sample sizes were doubled relative to the sample sizes per line described below. Approximately 3 days after collecting larva, we placed a half circle of Whatman #1 filter paper in each vial as a pupation substrate.

### Phenotypic assays

Here we briefly describe methods for each phenotypic measurement; full experimental details are available in the Supplemental Text. Pupation time was measured for three vials of 30 flies per line as the time in days for first instar larva to reach pupation. Fecundity was measured in 8 females per line by allowing single female flies to oviposit on grape agar for 48 hours and counting the number of eggs. In *D. melanogaster* and *Z. indianus,* diapause is characterized as the absence of vitellogenic oocytes in adult females and is assayed by exposing females to winter-like light and temperature conditions (Saunders et al. 1989; Lavagnino et al. 2020). We exposed 12 newly eclosed females per line to 21 days of 12°C, 10L:14D conditions, dissected their ovaries, and scored diapause as a binary trait using three cutoffs for ovary development (Erickson et al. 2020). Stage 8 is the beginning of yolk deposition in the oocyte; stage 10 includes a major developmental checkpoint and enlargement of the oocyte, and stage 14 is a mature egg ready for fertilization (King 1970; Soller et al. 1999; Mirth et al. 2019). For each cutoff, a fly was diapausing if no ovarioles had reached that stage and non-diapausing if at least one ovariole had reached that stage.

Morphological measurements were made for five males and five females from each line. We measured thorax length using a reticle on a stereomicroscope and dissected the right wing of each fly. We imaged the wing and used ImageJ v. 1.53k (Schneider et al. 2012) to fit an ellipse to the wing margins (Klaczko and Bitner-Mathe 1990; Bitner-Mathé and Klaczko 1999). We then extracted the major and minor axes of the ellipse and calculated the length of the major (*a*) and minor (*b*) radii. We calculated wing size as **√**(*ab*) (the geometric mean of the major and minor radii) and wing shape as *b/a*. Wing:thorax ratio was calculated as wing length (2*a*) divided by thorax length and is an inverse proxy for wing loading (Pétavy et al. 1997).

Chill coma recovery time (CCRT) was measured in approximately 6 males and 6 females from each line. We used Trikinetics Drosophila Activity Monitors to measure the time in seconds for flies to regain locomotion following a 2 hour exposure to 0 °C conditions. Four days later, we exposed the same flies to 45 minutes at −5 °C to measure the proportion surviving an acute freeze as freeze tolerance. Starvation tolerance was measured for 10-20 males and females for each line as the time in days that flies survived on 1% agar.

### Analysis: Post-colonization evolution

All analysis was conducted in R (version 4.3.3) using *data.table* (Dowle and Srinivasan 2019) for data manipulation and *ggplot2* (Wickham 2016) and *ggpubfigs* (Steenwyk and Rokas 2021) for plotting. We tested for post-colonization evolution by looking for changes in phenotype between the early and late season wild-derived flies without accompanying changes in the inbred control lines. We used linear mixed effect models implemented in *lme4* (Bates et al. 2015) to analyze the effects of capture season (early or late) on each continuous phenotype. We used generalized mixed effect models (GLMM) with a binomial error distribution for the binary traits of diapause and freeze survival. Fecundity (egg count) was zero-inflated, so we used a negative binomial mixed-effects model implemented in *glmmTMB* (Brooks et al. 2017). In most models, we included *control* status and *season* as binary variables and included a *season*control* interaction term in the model to test whether any changes in the controls were more or less extreme than changes in the wild-derived lines. We used the *emmeans* package (Lenth 2024) to calculate model-fitted least-square means and standard errors for plotting.

During initial analyses, we tested for an effect of *orchard* (Carter Mountain or Hanover Peach Orchard) as a fixed effect, but it was never significant after Bonferonni correction (P > 0.05), so we removed it from final models. *Sex* was always included as a fixed effect when traits were measured in both sexes. *Isofemale line* was always included as a random effect. Pupation and starvation tolerance data were collected from multiple replicate vials independently, and *vial* was included as a random effect nested under *isofemale line*. Although each experimental group (early, late, latitude) was assayed over two batches, we were unable to incorporate *batch* as a factor in most post-colonization models as batches were collinear with our temporal sampling and lacked the recommended number of levels to use as a random effect. *Batch* was included as a random effect for both the CCRT and freeze tolerance analysis due to the additional batches included in those experimental designs. Furthermore, many experiments were set up over the course of several days because adult flies did not all eclose at the same time, and we wanted flies to be approximately the same age at the time of analysis. For some phenotypes, we noticed that the phenotypes we gathered differed between flies set up earlier in the batch and those set up several days later. *Experiment date* was encoded as an integer variable starting with zero for the first day a given assay was set up within a batch. For assays started across multiple days (pupation time, diapause, and starvation tolerance), we initially included *experiment date* as a fixed continuous effect in the model and, if it was significant, included it as a random effect in the final model. A full description of models, including explanations for any exceptions to the analysis framework described above, can be found in the supplemental text. We determined significant effects of season and latitude using a Bonferroni correction for the twelve tests performed for each potential effect (α = 0.05/12 = 0.0042). Sexual dimorphism was tested for 7 phenotypes, with α = 0.05/7 = 0.007.

### Analysis: Lab evolution

We compared the phenotypes measured in the first experiment (after 3-4 generations of lab culture) and the second experiment (after 7-8 generations of lab culture) to test whether phenotypes had evolved in the lab for 12 isofemale lines. We included the control inbred lines in this analysis. All models were implemented with random and fixed effects as described above, including *assay generation* and *control* (as binary variables), and an *assay generation*control* interaction term, with *isofemale line* as a random effect.

### Analysis: Latitudinal variation

The effects of latitude were modeled with *latitude* of the collection locale as a fixed, continuous effect. We included *sex*, *isofemale line, vial, and experiment date* in the analyses as described above. Latitude experiments were conducted over two batches. In each analysis, we first tested for a batch effect by including *batch* as a fixed effect; if there was no significant effect, we removed it from the model. If there was an effect, we included *batch* in the final model as a fixed effect. Although we assayed inbred control lines alongside the latitude collections, they were not included in the models testing for an effect of latitude.

### Analysis: phenotypic correlations and principal components

To analyze relationships between phenotypes, we calculated the mean of each trait for each isofemale line (n = 174) used in the early, late, and latitude analyses (data from the second time point of the lab evolution experiment were excluded). We then used these line means to test for phenotypic correlations between traits and describe phenotypic change in multidimensional space. For correlation analysis, all phenotypes were compared in a Pearson correlation matrix; we only considered correlations significant if they passed a Bonferroni correction after testing n = 66 phenotype pairs (α = 0.05/66 = 0.00076). We plotted correlations using the *corrr* package (Kuhn et al. 2022).

For principal components analysis, we normalized the isofemale line means with z-scores for each phenotype. We used multiple imputation with the *MIPCA* method in the *missMDA* package (Josse and Husson 2016) to fill in lines with missing phenotypes. We then conducted PCA using the imputed phenotype matrix using the *prcomp* function. We used one-way ANOVAs and T-tests to test for significant differences in PCs between experimental groups.

As an alternative strategy to control for lab environment effects, we adjusted for the phenotypes of the controls prior to conducting correlation and PC analyses. We calculated the mean phenotype for the controls for each assay round (early, late, or latitude), and then subtracted that value from every isofemale line mean in that category. Therefore, each line mean became a measurement relative to the control. We then performed PCA and correlation analysis as described above using the adjusted line means.

## Results

### Post-colonization evolution

After accounting for differences in the inbred control line (lab effects) and correcting for multiple hypothesis testing, only wing size and wing:thorax ratio (an inverse estimate of wing loading) showed significant post-colonization evolution (Table 2, Figure 2, Figure 3). Wing size was significantly smaller in wild-derived late season flies (GLMM, P = 0.0007) and was slightly, but not significantly, larger in the inbred control lines assayed alongside the late season flies (P = 0.058), with a significant interaction between control status and season (P = 7.7 x 10^-5^). Wing:thorax ratio significantly decreased after colonization in wild-derived lines (P = 3.3 x 10^-7^), but did not differ in control lines (P = 0.128, Figure 3). Although not significant after Bonferoni correction, freeze survival increased later in the season (P = 0.015), and diapause at stage 14 decreased in the late season (P = 0.015) without significant changes in the controls (P > 0.57). Starvation time, pupation time, and wing shape all showed significant changes for post-colonization evolution (P < 0.0026 for all, Table 2), but the changes were in the same direction as significant changes in the inbred control line (P < 0.0011 for all, Table 2) with no interactions (P > 0.09 for all, Table 2), suggesting they were driven by inadvertent differences in rearing or assay conditions. See Table S1 for full results from all models.

**Figure 2:**
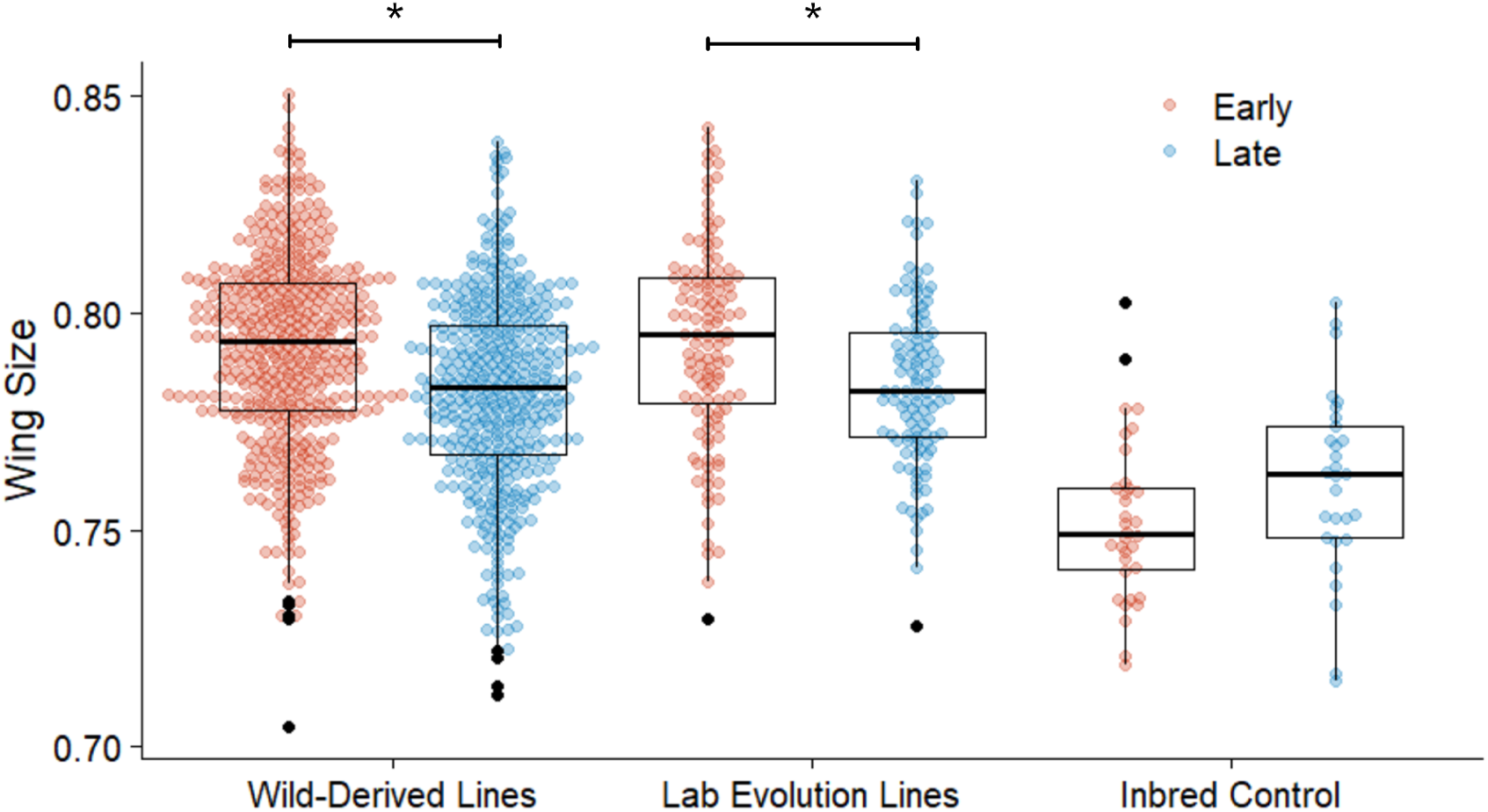
*Z. indianus* wing size evolves rapidly in the wild and in the lab. Wing size is the geometric mean of the major and minor radii of an ellipse fitted to the wing. Early and late describe when the flies were assayed; for the lab evolution experiment, the same lines were assayed at two different generations. Points illustrate all individuals measured (n = 1079 total); boxplots show median and quantiles. Asterisks indicate Bonferroni-corrected P < 0.05 in a mixed effects linear model (see Table S1 for full model results).

**Figure 3:**
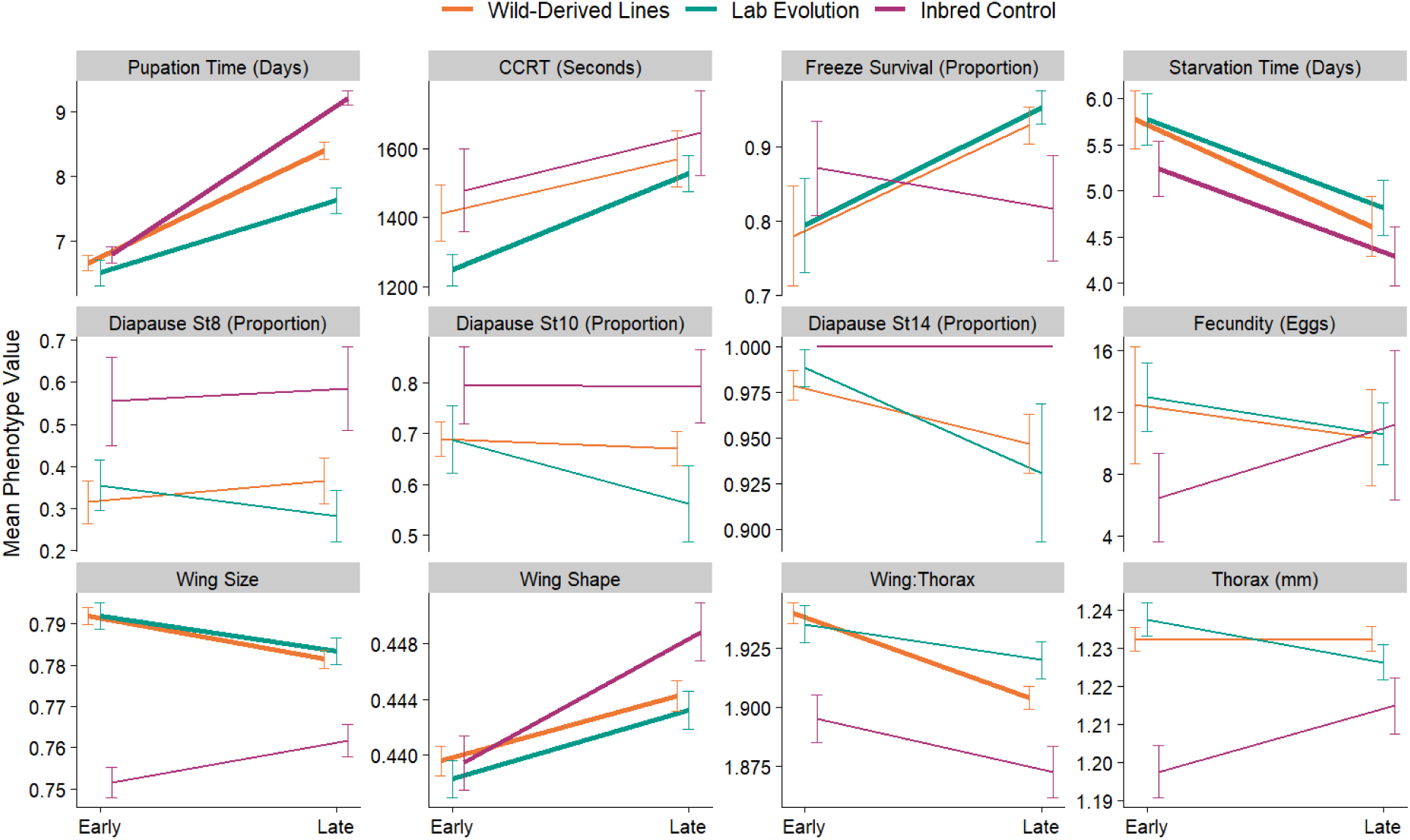
Post-colonization evolution, lab evolution, and lab environment effects for 12 phenotypes in *Z. indianus*. The endpoint of each line shows the least-squared model-fitted mean for each timepoint and error bars show model standard errors. Wild-derived lines (orange) were collected in August (early season) and November (late season) and measured for each phenotype after 3-4 generations of laboratory culture. Lab evolution lines (green) were collected and measured in the early season and remeasured alongside the late season flies after several additional generations in the lab. Inbred controls (purple) are a single inbred line that was assayed alongside all experimental flies. Bold lines indicate significant differences between early and late season phenotypes in mixed-effects linear models after Bonferroni correction (see Table 2; Table S1).

**Table 2:** P-values for main effects in mixed-effects linear models. Bold text indicates P-values that passed Bonferroni correction. Dashes indicate effects that were not assessed. See Table S1 and supplemental text for full model information and results.

|  | Temporal experiments |  |  |  |  |  | Latitude experiment |  |
| --- | --- | --- | --- | --- | --- | --- | --- | --- |
|  | *Lab environment (controls)* | *Post-colonization evolution (wild-derived)* | *Interaction (season: control)* | *Sex* | *Lab evolution (assay generation)* | *Interaction (assay generation: control)* | *Latitude* | *Sex* |
| pupation time | **<2.0x10<sup>-16</sup>** | **3.4x10<sup>-16</sup>** | 0.059 | — | **3.1x10<sup>-10</sup>** | **3.5x10<sup>-4</sup>** | 0.044 | — |
| CCRT | 0.339 | 0.212 | 0.7 | 0.406 | **7.85x10<sup>-5</sup>** | 0.546 | 0.864 | 0.316 |
| freeze survival | 0.57 | 0.015 | 0.152 | **3.5x10<sup>-4</sup>** | **3.4x10<sup>-4</sup>** | 0.008 | 0.818 | 0.042 |
| starvation time | **9.0x10<sup>-7</sup>** | **1.2x10<sup>-11</sup>** | 0.96 | **3.7x10<sup>-8</sup>** | **1.0x10<sup>-9</sup>** | 0.975 | 0.74 | 0.534 |
| fecundity | 0.374 | 0.663 | 0.264 | — | 0.392 | 0.027 | 0.817 | — |
| diapause st 8 | 0.819 | 0.327 | 0.887 | — | 0.334 | 0.816 | 0.245 | — |
| diapause st 10 | 0.979 | 0.687 | 0.539 | — | 0.136 | 0.239 | 0.047 | — |
| diapause st 14 | 1 | 0.015 | — | — | 0.021 | — | 0.563 | — |
| wing size | 0.059 | **7.57x10<sup>-4</sup>** | **7.73x10<sup>-5</sup>** | **<2.0x10<sup>-16</sup>** | **4.1x10<sup>-4</sup>** | **3.22x10<sup>-4</sup>** | 0.06 | **<2.0x10<sup>-16</sup>** |
| wing shape | **0.001** | **0.003** | 0.093 | 0.012 | **5.5x10<sup>-5</sup>** | 0.088 | 0.809 | **9.3x10<sup>-4</sup>** |
| wing:thorax | 0.128 | **3.37x10<sup>-7</sup>** | 0.375 | **5.84x10<sup>-11</sup>** | 0.037 | 0.613 | 0.053 | **<2.0x10<sup>-16</sup>** |
| thorax | 0.088 | 0.986 | 0.089 | 0.009 | 0.022 | 0.007 | 0.412 | 0.138 |

### Lab evolution

Chill coma recovery time and freeze survival both significantly increased with additional generations of lab rearing (Figure 3, P = 2.96 x 10^-6^ and P = 0.00023, respectively), and there was no accompanying change in the inbred control lines for either phenotype (P = 0.762, P = 0.57, respectively). Wing size significantly decreased with additional lab rearing (P = 0.00046, Figure 2, Figure 3) and did not differ significantly in the controls (P = 0.058). Starvation tolerance, pupation time, and wing shape all significantly changed between the two assay generations (P < 7.6 x 10^-5^ for all, Table 2, Figure 3), but these changes were accompanied by significant changes in the controls in the same direction (Table 2), as described in the previous section.

### Latitudinal variation

No phenotypes showed significant latitudinal clines after correcting for multiple hypothesis testing (Figure 4, Table 2), but some notable trends were observed. Wing size and diapause scored at stage 10 both increased with latitude (P = 0.06, P = 0.047, respectively), and time to pupation decreased with latitude (P = 0.039).

**Figure 4:**
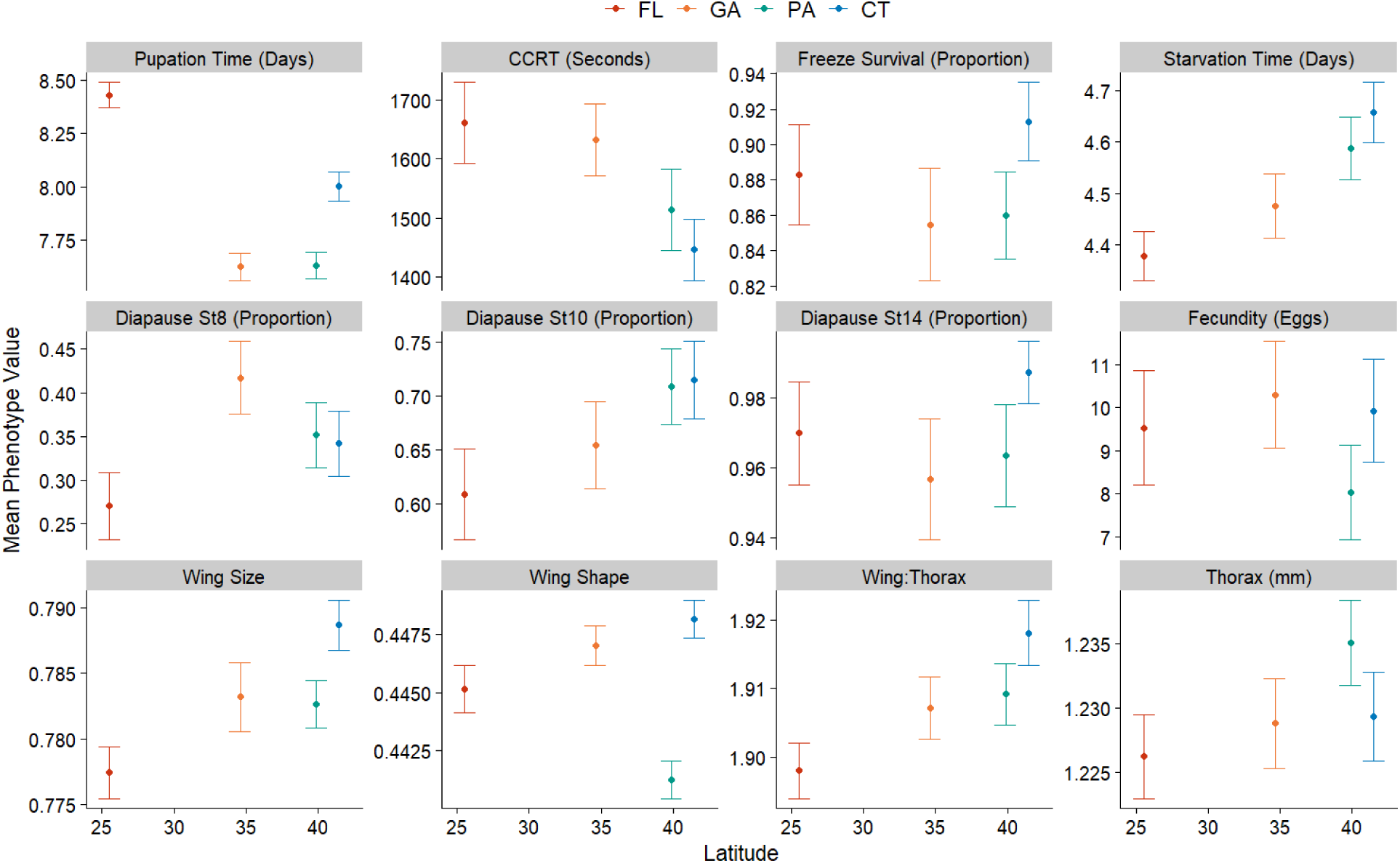
Limited latitudinal phenotypic variation in *Z. indianus*. Points represent the mean of all individuals from each population for a given phenotype, and error bars represent standard error of the mean. No phenotypes showed significant changes with latitude in the final models after Bonferroni correction.

### Sexual dimorphism

In both the latitude and post-colonization evolution experiments, we found significant sexual dimorphism for wing size and wing:thorax ratio after Bonferroni correction (Figure S2), with females having larger wings and higher wing:thorax ratios (GLMM, P < 5.84 x 10^-11^ for each analysis, Table 2). Additionally, in the temporal experiment only, we observed sexual dimorphism for starvation tolerance and freeze survival, with females having higher stress tolerance in both cases (P < 0.00035 for both, Table 2). In the latitude experiment, females had significantly broader wings (increased minor radius: major radius ratio, P = 0.00093). No significant sexual dimorphism was observed for chill coma recovery time or thorax size, though females tended to have larger thoraces in both experiments.

### Phenotypic correlations

When examining phenotypic correlations between all isofemale line means, we found significant correlations between 14 pairs of traits (Figure S3A), seven of which were from the same trait category (morphological traits or diapause). We suspected that some correlations might be driven by lab environment effects that produced large differences between the early and late seasons across multiple phenotypes (see Figure 3). We subtracted the control phenotype for each experiment from each isofemale line mean to adjust for potential lab effects. When we did so, we found only one significant correlation between traits from different categories (pupation time and diapause at stage 10 were negatively correlated, r = −0.266, P = 6.2 x 10^-4^, Figure S3B), though some diapause and morphological traits remained correlated to other measures in the same category.

### Multidimensional analysis

When accounting for variation in all 12 traits with principal components analysis (Figure 5A), we found that early and late populations significantly differed in both PC1, which explained 22.8% of variation (T-test, t = −5.18, df = 91.5, P = 1.3 x 10^-6^) and PC2, which explained 18.4% of variation (t = −6.2, df = 101.2, P = 1.2 x 10^-8^). We did not find significant temporal changes in PC3 or PC4 (t-test, P > 0.95 for both). Notably, the controls assayed alongside these groups had nearly identical values for PC2 but differed for PC1 (Figure 5A, triangles), suggesting that PC1 primarily describes lab environment effects and PC2 describes biological variation between the early and late samples. PC2 was strongly influenced by wing size, but most variables we measured (except fecundity and chill coma recovery time) correlated with both PC1 and PC2 (Figure 5C), suggesting this result reflects collective phenotypic changes and is not driven by a few traits. After subtracting the control means from each phenotype and recalculating PCs (Figure S4A), the early and late populations differed significantly for PC1 (t = 4, df = 81.4, P =0.0001), PC 2 (t = −5.27, df = 102.9, P = 7.6 x 10^−7^) and PC3 (t = −2.71, df = 102.5, P = 0.008), but not PC4 (P = 0.66).

**Figure 5:**
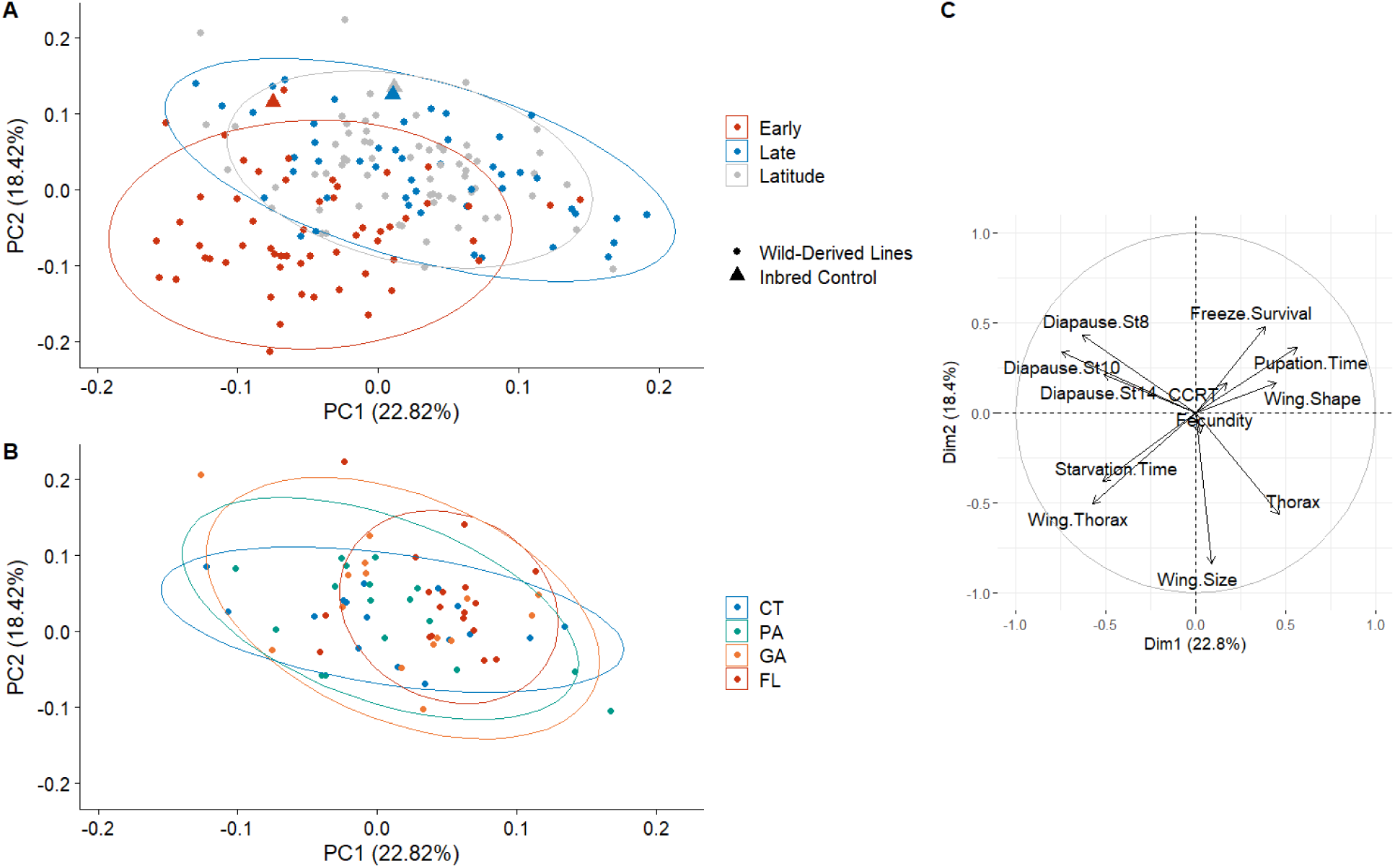
Principal component analysis of phenotypes. A) Principal components analysis for wild-derived flies caught early in the season, late in the season, and from four different latitudes (combined here for visual clarity). Each point represents one isofemale line. Triangles represent the inbred control line phenotyped alongside each group. Data ellipses were drawn using a multivariate t-distribution. B) PCA of the four latitudinal populations. C) Ordination plot for the 12 phenotypes used for the PCA; PCs were calculated using normalized line means for each phenotype from the combined seasonal and latitudinal experiments.

The four latitudinal populations did not differ significantly in PC1 through PC4 (ANOVA P > 0.25 for each, Figure 5B). However, after subtracting control values and recalculating PCs (Figure S4B), there was a slight difference between populations in PC2 (ANOVA, F = 2.97, df = 62, P =0.039) but not PC1, 3 or, 4 (P > 0.64). The latitudinal difference in PC2 was driven by a Florida-Connecticut difference (Tukey post-hoc test, P = 0.03), but there were no other significant differences between individual populations.

## Discussion

In this study, we tested for rapid evolution of fitness-related traits in invasive African fig flies (*Zaprionus indianus*) in the North American invaded range using rigorously controlled common-garden experiments involving over 15,000 flies from 174 isofemale lines. Looking at individual traits, we found that smaller wing size evolved following temperate colonization, resulting in increased wing loading, but we found no other significant post-colonization evolution or latitudinal variation. However, when examining all phenotypes together with a multidimensional analysis, we found evidence for significant post-colonization evolution and limited evidence for subtle latitudinal variation. Our findings demonstrate the possibility of rapid evolution in invasive species after only a few generations in a novel environment. We also found that several phenotypes evolved rapidly over just a few generations of lab rearing, and that measurements of many phenotypes are subject to the lab environment and batch effects despite our efforts to maintain carefully controlled experimental conditions. These findings emphasize the importance of accounting for potential lab evolution and assay variability when conducting any type of temporal phenotyping experiments for evolutionary studies.

### Rapid post-colonization evolution in Z. indianus

*Z. indianus* populations evolved smaller wings following colonization of two Virginia orchards during the 2022 growing season. As this decrease was not accompanied by a decrease in body size, this change produced an increase in wing loading (decreased wing:thorax ratio). We also observed a trend, not significant after multiple hypothesis testing, of larger wings at higher latitudes. Rapid changes in wing morphology are not unexpected in drosophilids; changes in wing shape that recapitulate a cline found in the native range were documented within a single decade following the invasion of *D. subobscura* in South America (Huey et al. 2000; Gilchrist et al. 2001). Aspects of wing morphology can change rapidly during a single growing season in both the temperate species *D. lutescens* and the cosmopolitan species *D. melanogaste*r (Önder and Aksoy 2022; Ueno et al. 2023). These changes can be favorable in different thermal environments; lines artificially selected under cold temperature evolved larger wings in *D. melanogaster* (Partridge et al. 1994) and wing size increases with latitude in both North American and South American *D. melanogaster* (Coyne and Beecham 1987; Zwaan et al. 2000). While the latitudinal trend we observed matches these findings, the finding of smaller wings later in the season as temperatures become cooler does not, suggesting other selective pressures may influence wing size in *Z. indianus* following colonization. For example, wing size is important for dispersal and range expansion in a variety of insects (Buckley et al. 2012; Renault 2020; Jahant-Miller et al. 2022). Although the mechanism of the yearly dispersal of *Z. indianus* through temperate regions of North America is unknown, larger wings may facilitate or enhance dispersal such that early colonizing populations are likely to be large-winged. Following colonization by larger-winged individuals, energetic tradeoffs in the new environment may favor smaller wings and investment in other traits despite increased wing loads. For example, larger-winged *D. melanogaster* living at high elevation in Africa lay fewer eggs (Lack et al. 2016), presumably due to tradeoffs. However, we did not identify any non-morphological traits that were correlated with wing size in our analysis, so the source of any potential tradeoff remains to be determined.

Wing size evolution has been studied in *Z. indianus*, but not in North American populations. Despite a bottleneck from invasion, *Z. indianus* retains substantial wing morphology variation in South America (Loh 2005). Wing size was larger in subtropical populations in South America than in equatorial African populations of *Z. indianus* less than a decade following invasion (Loh et al. 2008) and this finding was replicated in a second study (Lavagnino et al. 2020b). Our latitudinal trend of increasing wing size at higher latitudes agrees with previous studies of *Z. indianus*: David et al. (2006) identified subtle wing size clines with wing size increasing with latitude in both Africa and India, in agreement with previous studies of Indian populations (Karan et al. 1999). However, the same study found no latitudinal trends in wing size in invasive South American populations, so our study is the first to document a potential emerging cline in the invaded range in the Americas. In addition to genetically determined differences in morphology, the plasticity of wing size and shape is more extreme in subtropical populations relative to equatorial populations, suggesting that wing size and shape variation may be an important component of coping with local environmental conditions in this species (Loh et al. 2008). Given known variation in wing morphology, the rapid evolution of wing size we observed in *Z. indianus* is not unexpected and emphasizes the species’ capacity for rapid evolution.

Aside from wing size (and wing:thorax ratio, which is influenced by wing size), no other individual traits showed significant post-colonization evolution. The proportion of flies surviving an acute freeze increased late in the season, though this difference was not significant after correcting for multiple hypothesis testing. Temperatures dropped to near freezing in both Charlottesville, VA, and Richmond, VA approximately two weeks prior to our late season collection in mid-November (data viewed on wunderground.com); this event may have selected for *Z. indianus* slightly more tolerant of acute freezes late in the season. Drosophilids are known to evolve rapidly when selected in thermally varying environments (MacMillan et al. 2009; Tobler et al. 2015; Gerken et al. 2016). Chill coma recovery time, a more commonly measured proxy for thermal tolerance (Andersen et al. 2015) showed no temporal variation in our data and had little influence on the PCA, suggesting it does not vary with seasonal environments in *Z. indianus.* This finding differs from the cosmopolitan species *D. melanogaster*, which shows seasonal changes in its response to thermal stress (Behrman et al. 2015). Diapause incidence also evolves seasonally in *D. melanogaste*r (Schmidt and Conde 2006; Erickson et al. 2020) but did not change in *Z. indianus*. Collectively, non-morphological phenotypes that are known to evolve on short seasonal timescales from other drosophilid systems did not evolve following colonization in *Z. indianus*. Assuming an ∼18 day generation time at 22 °C (Nava et al. 2007), Virginia populations of *Z. indianus* may experience 4-5 generations in the three months between our early and late sampling, which is fewer than the ∼10 generations *D. melanogaster* experiences during their longer growing season (Bergland et al. 2014). Fewer generations and a lack of overwintering selection may limit the extent of evolution that occurs following a single colonization event.

However, despite the relative lack of post-colonization changes in individual traits, we found evidence for evolution across traits in multidimensional space. Multidimensional traits are thought to be important measures of fitness for organisms in changing environments (Laughlin and Messier 2015). While temporal differences in PC1 can likely be explained due to laboratory effects, as evidenced by shifts in the PC1 values in the controls, PC2 changed significantly between the early and late populations in the wild-derived but not control flies. Most of the traits we measured loaded on PC2, suggesting that the majority of traits contributed to this subtle population shift, which held up even after adjusting phenotypes for laboratory effects observed in the controls. So, although the differences in most individual traits were not significant, collective subtle changes in multiple traits might change the fitness of *Z. indianus* when colonizing temperate environments, fueling their successful invasion.

### Reproductive traits and environmental conditions in Z. indianus

We noticed several interesting patterns when examining data related to reproduction in *Z. indianus*. First, though our sample size is limited, isofemale success rates were lower later in the season and at higher latitudes (Table 1). Although an imperfect measure of reproduction, these patterns suggest previous environmental exposure of wild-captured flies influenced their egg laying in the lab. We found no differences in fecundity across any of the common garden experiments, suggesting that the field-based differences in fecundity were likely plastic responses to environmental conditions prior to collection. *Z. indianus* males are sterile (Araripe et al. 2004) and females enter diapause (Lavagnino et al. 2020a) when reared at low temperatures, which might explain the lower isofemale success rates seen in later months and higher latitudes. Alternatively, low population densities in the field (Rakes et al. 2023) might have limited the number of females that successfully mated prior to capture, given that *Z. indianus* are sperm-limited (Gleason et al. 2024). Although a previous study found a latitudinal cline in ovariole number in Indian populations of *Z. indianus* (Karan et al. 2000), any potential variation in reproductive investment did not translate to fecundity differences in our study. Interestingly, we found that ∼30% of lab-reared females laid zero eggs, despite multiple days of access to mates and food prior to the start of the experiment, suggesting that some other biological factor influences fecundity in this species. The low fecundity rates of both field-caught and lab-reared flies are puzzling given the overall success of *Z. indianus* as an invasive species. The causes of low fecundity rates and the alternative ecological mechanisms that may contribute to *Z. indianus*’ rapid population growth in newly invaded environments (Rakes et al. 2023) warrant future study.

Though the trend was not significant, we found that rates of diapause subtly increased with latitude in *Z. indianus*. This result is somewhat surprising because it matches patterns observed in *D. melanogaster*, which is thought to overwinter, unlike *Z. indianus* (Schmidt et al. 2005; Schmidt and Paaby 2008). In higher-latitude environments with harsher winters, diapausing genotypes are more likely to survive winter and increase in frequency. Since *Z. indianus* likely does not overwinter (Pfeiffer et al. 2019) and our latitudinal samples were collected in October, before the onset of conditions that would likely favor diapausing genotypes, the cause of this trend is unknown and could be due to another trait that is correlated with diapause or trades off with diapause. We also did not identify a tradeoff between ability to diapause and fecundity, as might be predicted from findings in *D. melanogaster* (Schmidt and Conde 2006). Collectively, our results suggest that the reproductive biology of female *Z. indianus* is influenced by the environment, and further understanding this species’ reproduction may be important to controlling it in agricultural settings.

### A lack of predictable latitudinal variation in Z. indianus

Aside from the wing size and diapause trends already discussed, we found little evidence of latitudinal clines in *Z. indianus* phenotypes, despite abundant data on latitudinal clines in similar traits in other drosophilids (Coyne and Beecham 1987; Azevedo et al. 1998; Zwaan et al. 2000; Hoffmann et al. 2002; Ayrinhac et al. 2004; Fabian et al. 2015; Rohner et al. 2018). We observed a trend that Florida populations developed more slowly than northern populations. South American populations of *Z. indianus* develop more slowly than those from Africa (Lavagnino et al. 2020b); our data suggest that this trend may be reversed as flies move northward. Starvation tolerance also decreases with latitude in Indian populations of *Z. indianus* (Karan et al. 1998). One possible explanation for the lack of latitudinal patterns in North America is that a single growing season offers insufficient time for substantial latitudinal clines to evolve. All our samples were collected within several weeks of each other in October 2022; given a northwards recolonization model, the northernmost populations may have been locally established for a relatively short time prior to collection. In *D. melanogaster*, many clinal traits are thought to be the result of selection imposed on overwintering populations, with harsher winters at more northern latitudes (Flatt 2020). Since *Z. indianus* is apparently not able to overwinter and instead reestablishes each year (Pfeiffer et al. 2019; Rakes et al. 2023), traits favorable for overwintering would not be selected at higher latitudes. However, our collective findings suggest a high degree of phenotypic variation in *Z. indianus* and the potential for rapid evolution. As northern winters become warmer with climate change (Marshall et al. 2020), it is possible that *Z. indianus* populations may become permanently established at higher latitudes, which would create more potential for latitudinal clines to evolve. We also note that several phenotypes shown in Figure 4 appear to show latitudinal trends, but some of these trends were driven by batch effects and were not significant in the final models after including all predictor variables. Lastly, some traits (for example, wing shape) show substantial variation that does not fit a linear trend; bottlenecks from small founding populations upon recolonization could potentially produce populations with phenotypes that diverge but lack latitudinal patterns.

### The potential for laboratory adaptation in isofemale lines

We observed several instances of phenotypic changes in isofemale lines in the lab, with trait measurements changing significantly following 3-4 additional generations of lab rearing. Interestingly, two traits that showed significant changes were both related to cold tolerance, but they evolved in opposite directions. Chill coma recovery time became longer, suggesting lab-adapted flies were less tolerant of a mild cold stress. However, the proportion of flies surviving an acute freeze increased. A meta-analysis found that *Drosophila* tend to evolve both negative and positive changes to stress tolerance in the lab, but most substantial changes are negative, meaning lab reared populations become less stress tolerant (Hoffmann and Ross 2018). Our result is surprising because *D. melanogaster* selected for improved chill coma recovery time also showed a correlated increase in tolerance of acute cold stress (Anderson et al. 2005). However, as *Z. indianus* likely has not historically experienced selection for survival under cold conditions (Pfeiffer et al. 2019), the biology of its cold responses may be different. This result could alternatively be related to our experimental design: we re-used the same flies for CCRT and freeze survival assays. It is possible that some flies died due to the stress of the CCRT experiment, and the CCRT may have served as a cold hardening event for the flies used in the freeze experiment (Czajka and Lee 1990). Therefore, our freeze experiment may in part be measuring cold hardening rather than baseline cold tolerance and may have been altered by lab rearing. Regardless of the individual traits that changed in the lab or their directions, the finding of trait evolution in 3-4 generations of lab rearing represents an important finding for *Z. indianus* invasion biology as it demonstrates the species’ ability to rapidly evolve in new environments, potentially driving future range expansion.

### Relevance to evolutionary studies

Despite our attempts to carefully control experimental conditions over an experiment that spanned several months, we found strong evidence of uncontrolled variation in lab conditions that altered phenotypes. Several phenotypes showed parallel changes in the inbred control and wild-derived lines assayed at different times, strongly suggesting the variation was due to uncontrolled laboratory factors since our inbred line should be genetically uniform and phenotypically consistent. This finding is similar to those of Stone et al. (2020), who found that a lab-reared population and field-reared population showed similar temporal changes in cold-hardening, suggesting that the variation was caused by assay variation rather than selection in the field. This finding is of great relevance to studies of rapid evolution. Studies that assay temporal variation in flies are common in *Drosophila* literature and have documented a variety of phenotypic changes on short time scales (Schmidt and Conde 2006; Behrman et al. 2015, 2018; Aggarwal et al. 2021; Grainger et al. 2021; Rudman et al. 2022). Most of these studies limited the number of generations that flies were held in the lab to minimize lab adaptation, opening the possibility for unintentional variation in rearing or assay conditions over time. These studies may have had less lab-induced variation than we observed, but without controls it is impossible to know for certain. We recommend that future studies of temporal changes in fly populations use control populations as a reference when possible. This finding is also relevant for longitudinal sampling experiments (Eccard and Herde 2013; Mangan et al. 2022), multigenerational experiments (Foucault et al. 2018; Toyota et al. 2019), or experimental evolution studies that compare endpoints to starting phenotypes (Kawecki et al. 2012). While not every experimental design or organism allows for control lines or reference samples like the ones used here, the potential for assay variation should generally be taken into consideration for temporal studies.

### Conclusions and future directions

With the exception of wing size, we found limited evidence for heritable changes in *Z. indianus* phenotypes across space or short time scales following invasion. While the evolution of smaller wings may provide a fitness advantage following colonization, this finding suggests that additional factors not measured here are responsible for the enormous success of *Z. indianus* in the invaded range. First, *Z. indianus* likely has ecological advantages not captured in the laboratory environment; for example, it is an unfavorable host for North American parasitoids, allowing it to outcompete co-occurring drosophilids in the presence of parasitoids (Walsh-Antzak and Erickson 2024). Additionally, *Z. indianus* may be inherently tolerant of a wide range of conditions, as evidenced by its large geographic range in Africa and broad thermal niche (Comeault et al. 2020) and its ability to use a wide variety of host fruits (Yassin and David 2010). Third, it might possess a high degree of phenotypic plasticity, which could enable success in a wide variety of environments (Yassin et al. 2009). Lastly, rapid adaptation involving fitness-relevant traits beyond those measured here, such as behavior, male reproductive traits, or immune function, may occur in novel environments. Beyond the biology of this fascinating invasion, our results highlight the importance of rigorous design when conducting temporal common garden experiments. We found significant differences in several phenotypes when we assayed an inbred control line at multiple timepoints, suggesting variation in the laboratory environment contributed to those differences. Future studies will be required to determine the extent to which adaptive changes contribute to *Z. indianus*’s fitness in the wild, and whether the changes we observed here are repeated in other colonization years. Longer-term studies might identify how North American populations are diverging from intermediate South American and ancestral African populations and whether milder winters might permit the evolution of permanently established temperate populations, which could dramatically increase the pest potential of *Z. indianus*.

## Data Availability Statement

All raw phenotyping data used for analysis has been deposited on Dryad at: https://doi.org/10.5061/dryad.gqnk98sxp. Code used for analysis and plotting is available on Zenodo via the above link, and a backup copy can be found on Github: https://github.com/ericksonp/Zindianus_phenotyping_2022

## Author contributions

Conceptualization: PE

Data curation: WG, LR

Formal analysis: WG, LR, PE

Investigation: WG, LR, CC, AG, JH, SM, CW, JY, PE

Supervision: PE

Visualization: WG, PE

Writing-original draft: PE, WG, LR

Writing-reviewing and editing: PE, WG, LR, CW, JY

## Acknowledgments

The authors thank the owners and managers of the orchards and parks we visited for permission to collect flies. We thank Alan Bergland and members of the Bergland lab for helpful feedback on this project.

