## Supplemental for "Rapid wing size evolution in African fig flies (*Zaprionus indianus*) following temperate colonization"

### Supplemental Text:

Below are detailed experimental procedures for the measurement of each phenotype in isofemale lines.

*Pupation time:*

Three vials of each line were monitored for pupation starting five days after larva collection. At the same time each morning, we counted the number of pupae in each vial. This procedure was repeated daily until no larva were visible and no additional pupae had formed for two consecutive days. Pupation times reported are the time from larva collection to pupation. Preliminary experiments in which we recorded pupation time and the number of adults to emerge each day revealed that pupation time and adult emergence time are highly correlated in *Z. indianus* isofemale lines (Figure S1, linear model, $R^2 = 0.95$, $P < 2 \times 10^{-16}$), so we did not record adult emergence times.

*Diapause:*

Adult *Z. indianus* began to eclose approximately 10 days after larva transfer (12 days after egg laying). For each line, 12 females were collected within 2 hours of eclosion. These females were placed in an incubator at 12°C with a 10L:14D light cycle in a vial of cornmeal-molasses food for three weeks. Flies were transferred to fresh food after 10 days. After a total of 21 days, females were frozen at -80°C until dissection.

We briefly immersed each fly in ethanol and then dissected the ovaries in a drop of phosphate buffered saline beneath a stereomicroscope with transillumination. For each fly, we recorded the stage of the most advanced ovariole from 6-14 (King 1970) and used the cutoffs described in the main text to determine if each fly was in diapause.

*Fecundity:*

We placed approximately 16 females and 8 males in a vial together for four days prior to the assay to allow time for mating. We assayed fecundity in 24-well tissue culture plates containing 1 mL of 3% agar, 30% grape juice concentrate per well. For each line, we placed eight anesthetized females into individual wells of the plate. We secured flies with a layer of soft medical foam padding between the wells and the lid. We incubated the plates for 48 hours and then removed the adult flies and counted the number of eggs and larvae in each well under a dissecting microscope. The males and extra females not used for the assay were frozen at -20°C for morphological measurements.

*Wing and body size:*

We collected morphological measurements on five males and five females from each line. To measure thorax length, we viewed the fly laterally under a stereomicroscope and measuring a straight line from the anterior tip to the posterior tip of the notum using a reticle. We removed the right wing from each fly, mounted it between two coverslips, and imaged it on a stereomicroscope outfitted with a digital camera. We used ImageJ to create a modified implementation of the previously described ellipse fitting method for wing morphology (Klaczko and Bitner-Mathe 1990; Bitner-Mathé and Klaczko 1999), which has been used to describe wing morphology in *Z. indianus* (Loh and Bitner-Mathé 2005; Loh et al. 2008). We drew an ellipse with one vertex at the distal end of the L3 vein. We adjusted the width and height of the ellipse to match the curvature of the wing such that the ellipse passed through the distal ends of L2, L4, and L5, and the proximal junction of veins L4 and L5.

*Chill coma recovery:*

We measured chill coma recovery time (CCRT) in approximately 12 individuals (6 of each sex) from each line. We placed individual flies in 5 mm *Drosophila* Activity Monitor (Trikinetics, Waltham, MA) tubes containing ~1 cm of cornmeal molasses medium and allowed them to recover overnight. Immediately prior to the assay, we pushed the cotton closure down so that flies had approximately 1 cm of space to move within the tube. We placed the tubes vertically in an ice bucket and put the ice bucket in a freezer set to 0°C. After two hours, the tubes were removed from the ice and placed in Drosophila Activity Monitors so that the fly was resting on the cotton, immediately in front of the infrared beam. This setup ensured that the first steps taken by the fly would be recorded as a break in the infrared beam. We left the monitors undisturbed until all flies regained locomotion. We recorded CCRT as the time in seconds between removing the flies from ice and the first movement. Each experimental batch was split into two sub-batches for the CCRT experiment; we accounted for these batches in models when possible. The flies were held in the same activity monitor tubes for the freeze tolerance experiment.

*Freeze tolerance:*

Four days after the chill coma recovery experiment, we removed any tubes containing dead flies. Tubes with surviving flies were placed in a freezer set to -5 °C for 45 minutes for a cold shock treatment. We allowed the flies to recover for 24 hours, then scored each fly as alive or dead based on the ability to right itself after agitation.

*Starvation tolerance*

We measured starvation tolerance of each line by placing ten flies of the same sex in a vial containing 10 mL 1% agar. We assayed one or two vials of each sex for each line depending on availability of adult flies. All flies were at least three days and no more than seven days post-eclosion at the start of the assay. We held the vials at 27°C and counted the number of survivors at the same time every day until all flies died.

*Model fitting:*

Below are the final generalized, mixed-effect linear models used for each phenotype in each experiment (* indicates binary phenotype with binomial GLM; †indicates zero-inflated model).

*Post-colonization evolution*

Wing size, Wing shape, Wing Load, Thorax

phenotype ~ season*control + sex + (1|line)

Chill coma recovery time, freeze survival*

phenotype ~ season*control + sex + (1|batch/line)

Fecundity†

phenotype ~ season*control + (1|batch/line)

Diapause (stage 8 and 10)*

phenotype ~ season*control + (1|experiment_date/line)

Diapause (stage 14)*

phenotype ~ season + (1|line)

*Note:* control *was removed when analyzing diapause at stage 14 due to high levels of multicollinearity between* control *and the* season*control *interaction.*

Starvation time

*phenotype ~ season*control + sex + (1|line/vial) + (1|experiiment_date)*

Pupation time

*phenotype ~ season*control + (1|line/vial)*

*Lab evolution*

Wing size, Wing shape, Wing Load, Thorax

*phenotype ~ assay_generation*control + sex + (1|line)*

Chill coma recovery time

*phenotype ~ assay_generation*control + sex*

*Note: line was not included as a random effect in this model due to singularity. Batch was not included due to low sample size across batches.*

Freeze survival*

*phenotype ~ assay_generation*control + sex + (1|line)*

Fecundity†

*phenotype ~ assay_generation*control + (1|line)*

Diapause (stage 8 and 10)

*phenotype ~ assay_generation*control + (1|experiment_date/line)*

Diapause (stage 14)

*phenotype ~ assay_generation + (1|line)*

Starvation

*phenotype ~ assay_generation*control + sex + (1|line/vial) + (1|experiment_date)*

Pupation

*phenotype ~ assay_generation*control + (1|line/vial)*

*Latitude evolution*

Wing size, Wing shape, Wing Load, Thorax

*phenotype ~ latitude + sex + (1|line)*

Chill coma recovery time, freeze survival*

*phenotype ~ latitude + sex + batch + (1|line)*

Fecundity[†]

*phenotype ~ latitude + (1|line)*

Diapause (stage 8 and 10)*

*phenotype ~ latitude + batch + (1|line)*

Diapause (stage 14)*

*phenotype ~ latitude +(1|line)*

Starvation

*phenotype ~ latitude + sex + batch + (1|line/vial) + (1|experiment_date)*

Pupation

*phenotype ~ latitude + batch + (1|line/vial) + (1|experiment_date)*

**Supplemental Table:**

**Table S1 (Excel File).** Linear model results from post-colonization, lab evolution, and latitude experiments. File is organized with a tab for each phenotype, with the results of linear models for each experiment (post-colonization, lab evolution, and latitude) on one page.

**Supplemental Figures:**

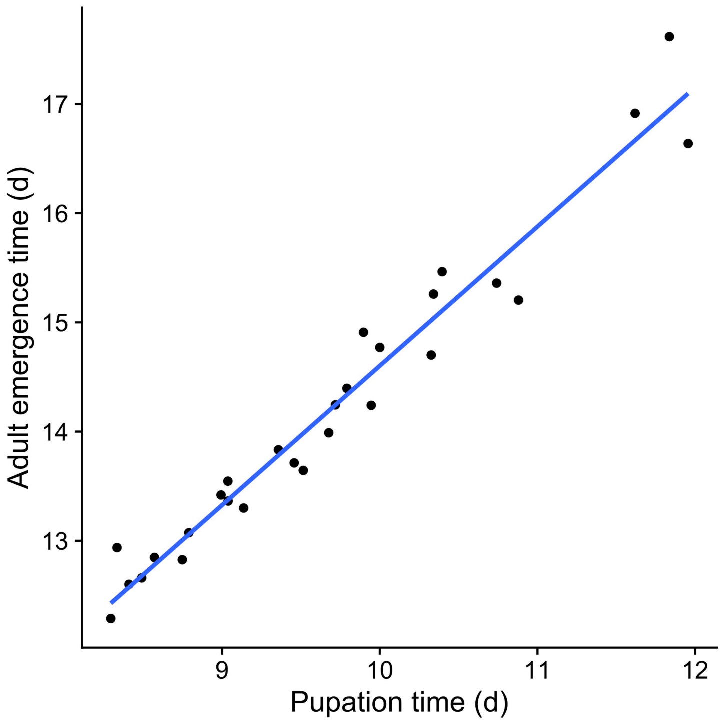


**Figure S1: Pupation time and adult emergence time are correlated in *Z. indianus*.** Each point represents the mean pupation and emergence time for individuals from a single isofemale line (n = 28). A mean of n = 110 flies were counted for each line. The isofemale lines used for this analysis were collected in 2019-2021 and were not part of the main phenotyping experiment.

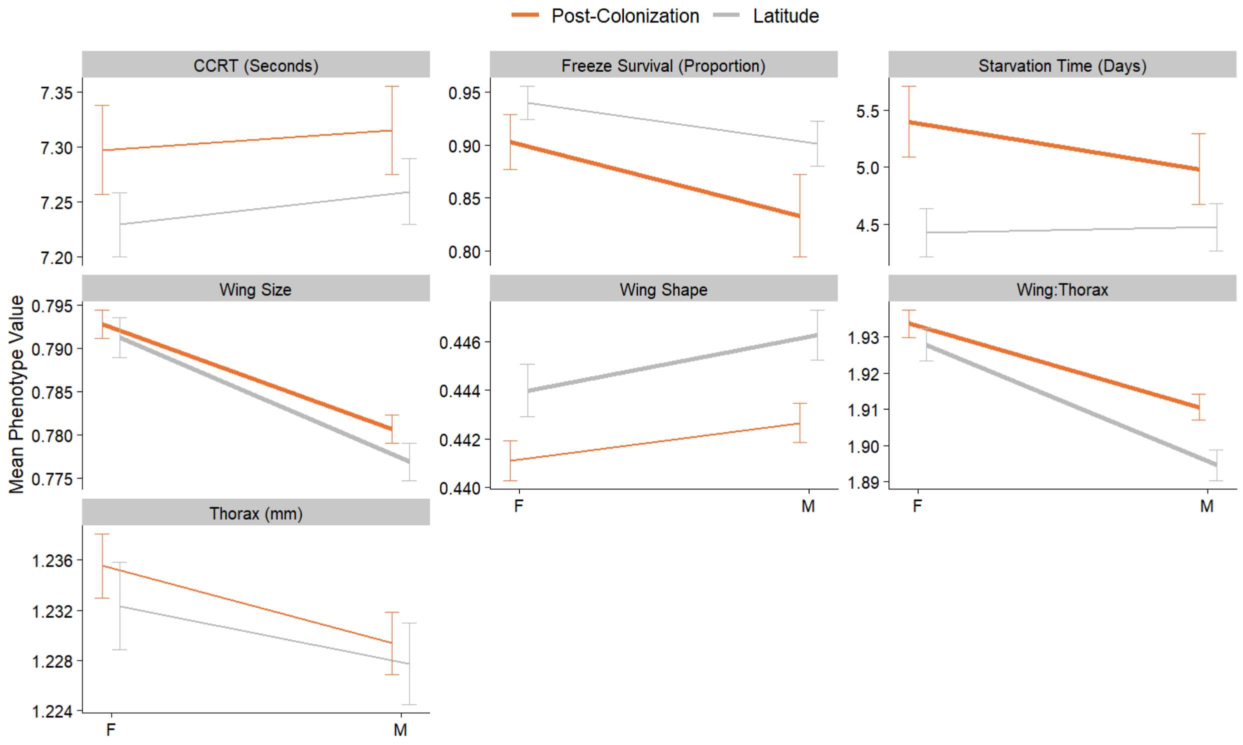


**Figure S2: Sexual dimorphism in *Z. indianus*.** Only phenotypes measured separately for the two sexes are shown. The end of each line shows the model-fitted least-square mean for each sex and error bars show standard errors. Data are shown separately for the post-colonization and latitude experiments. Bold lines indicate significant sexual dimorphism in mixed effects models following Bonferroni correction.

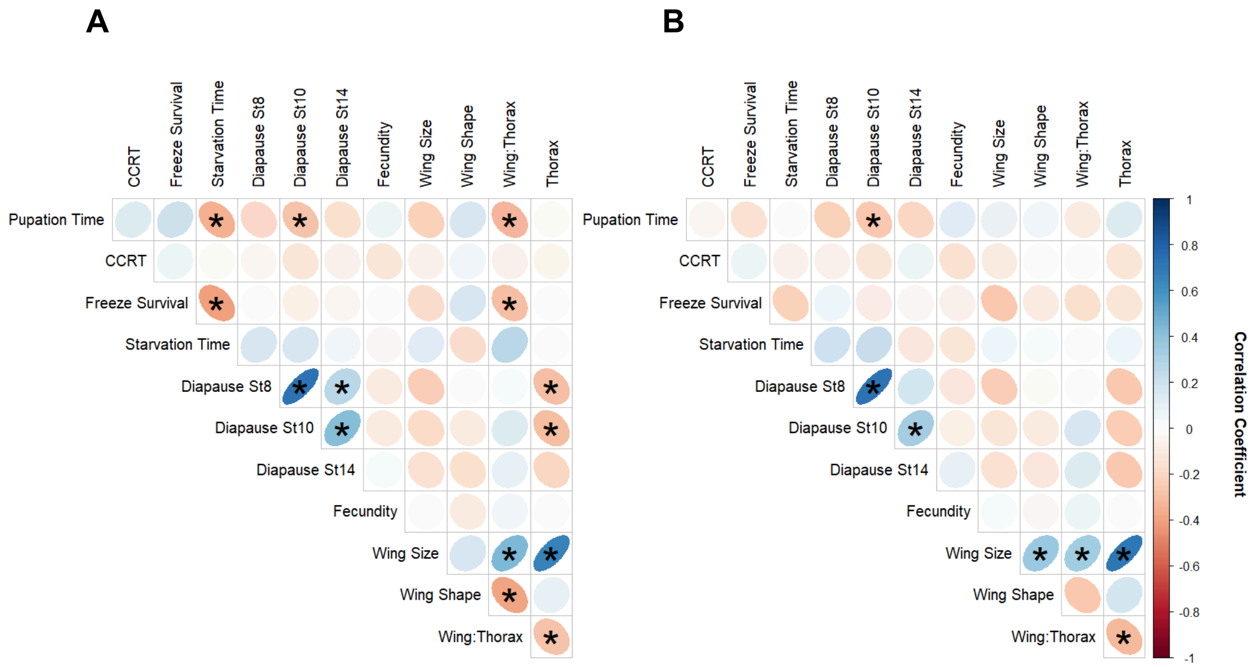


**Figure S3: Phenotypic correlations.** The color and size of the circle indicates the magnitude of the correlation coefficient; correlations significant after Bonferroni correction (n = 66 tests) are shown with asterisks. Line means for each phenotype were used for all correlations. (A) shows all raw phenotype correlations. In (B), the mean phenotype of the control line for each phenotyping group (early, late, or latitude) was subtracted from each line mean to remove environmental effects.

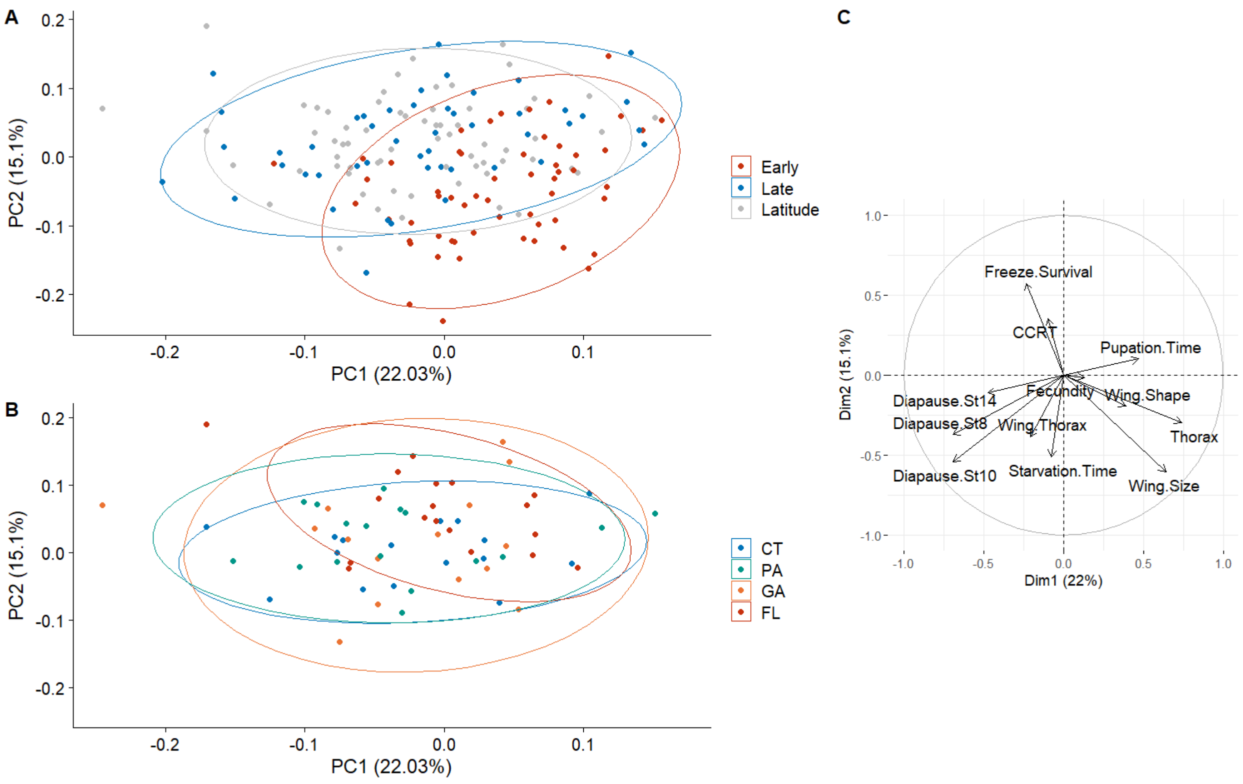


**Figure S4: Principal components analysis of phenotypes after adjusting for lab environment effects.** The mean of the control line for each phenotyping batch was subtracted from each isofemale line mean to account for lab variation. A) PCA of all data from the post-colonization and latitude experiment. B) PCA of four latitudinal populations. C) PCA loadings.
